# Co-Evolving Dynamics and Topology in a Coupled Oscillator Model of Resting Brain Function

**DOI:** 10.1101/2023.01.31.526514

**Authors:** Maria Pope, Caio Seguin, Thomas F. Varley, Joshua Faskowitz, Olaf Sporns

## Abstract

Dynamic models of ongoing BOLD fMRI brain dynamics and models of communication strategies have been two important approaches to understanding how brain network structure constrains function. However, dynamic models have yet to widely incorporate one of the most important insights from communication models: the brain may not use all of its connections in the same way or at the same time. Here we present a variation of a phase delayed Kuramoto coupled oscillator model that dynamically limits communication between nodes on each time step. An active subgraph of the empirically derived anatomical brain network is chosen in accordance with the local dynamic state on every time step, thus coupling dynamics and network structure in a novel way. We analyze this model with respect to its fit to empirical time-averaged functional connectivity, finding that it significantly outperforms standard Kuramoto models with phase delays. We also perform analyses on the novel structural edge time series it produces, demonstrating a slowly evolving topology moving through intermittent episodes of integration and segregation. We hope to demonstrate that the exploration of novel modeling mechanisms and the investigation of dynamics *of* networks in addition to dynamics *on* networks may advance our understanding of the relationship between brain structure and function.

## I. INTRODUCTION

Brain structure shapes function through dynamics. In neuroimaging, there have been many efforts to model the dynamics that link the brain’s anatomical network to the functional dependencies between regions as measured with BOLD fMRI [1–6]. A traditional approach uses dynamical systems modeling to capture the interactions between neural populations or brain regions [2, 6, 7]. Examples of such models range from biologically realistic implementations of integrate-and-fire neurons or neural masses [8] to simpler, more abstract, realizations of coupled oscillators, e.g. the Kuramoto model [9–12]. By allowing moment-to-moment dynamics to unfold on a fixed structural (often empirical) network, these models produce a time series of simulated neural activity that can be directly compared to empirical data.

An alternative approach to modeling brain function is by modeling network communication [13–15]. This class of models uses concepts from graph theory and network science to describe patterns of neural signaling, considering edges with respect to the role they may play in signaling strategies. Two broad families of brain network communication models have been explored in literature: routing protocols (communication via efficient, selectively accessed paths, e.g., the shortest topological path between two nodes—and diffusion processes) broadcasting along multiple network fronts or random walk dyanmics [16, 17]. The contrast between these two signaling strategies illustrates an important trade-off. On the one hand, routing via shortest paths is efficient with respect to transmission delays and energetic expenditure [18]. On the other, the use of shortest paths presupposes that nodes have global knowledge of the network beyond their immediate vicinity [14, 19]. Conversely, while simultaneous broadcast along every edge does not mandate these assumptions, signal retransmission under these models would require exceedingly large metabolic demands [18]. We argue that most implementations of dynamical models are akin to network communication via broadcasting, as their operation on a static network requires that all edges carry influence between their incident nodes at all time points. As such, models of coupled differential equations have thus far been unable to capture the wide array of possible signaling strategies described in the brain network communication literature.

We seek to take a step toward integrating the key strengths of both of these approaches by proposing a communication-inspired variation of the Kuramoto-Sakaguchi (K-S) model. The K-S model is a well-studied coupled oscillator model incorporating phase delays [9, 20], and has been used to capture BOLD [7, 10, 21–23], MEG [24], and EEG [25, 26] dynamics. Individual oscillators are set at an intrinsic frequency of 40-60 Hz, which can be understood to replicate locally synchronous neural firing in the gamma range. [10–12, 27]. The model’s phase-amplitude time series can be further processed to approximate low-frequency BOLD signal fluctuations, either by using a convolution with a haemodynamic response function or by implementing a dynamic model of the neurovascular response [22]. In certain coupling regimes, the model produces metastable behavior reminiscent of empirical brain activity [28–30] and calculation of the modeled functional connectivity matrix produces significant correlation to empirical FC [7, 10, 31].

When the Kuramoto-Sakaguchi model is used according to this paradigm, an oscillator is assigned to each of *N* brain regions (nodes). At each time step the phase, *θ*, of every oscillator is updated according to its intrinsic frequency, *f*, and the influence from other oscillators along a structural network (from empirical data) according to the equation:

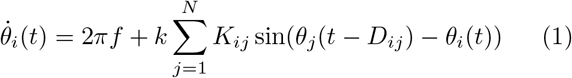

where *K* is a matrix that stores the weight of the structural connection between nodes *i* and *j*, *D_ij_* is the phase delay between nodes *i* and *j*, and *k* is a global coupling constant. The first term of this equation captures the intrinsic fluctuations of each oscillator, while the second term of the equation can be thought of as the “influence term”, which calculates the interactions between oscillators. A careful look at the influence term reveals that the influence is summed over all neighbors of each node (and hence over every edge) on each time step. The only situation in which a node does not contribute to its neighbor is the (exceedingly) rare instance that the sine of the difference of their phases is exactly equal to zero. The summation of influence over all edges implements a broadcasting communication process in which all edges are ‘active’ over the entire length of the simulation. The extent to which this form of dynamic communication may limit the predictive power of the model has yet to be considered.

Here we introduce a model that uses a subset of ‘active’ edges chosen in accord with the local dynamic state of the oscillators. On every time step each node selects a number of active edges according to local edge selection rules. Limiting nodal communication in this way has the added effect of coupling time-varying structure and function. Interactions between nodes are determined by the selected edges which then influence the selection of active edges on the next step. Structure thus shapes dynamics, and dynamics selects structure in turn, in an ongoing reciprocal ‘co-evolutionary’ dialogue. Selecting a limited number of active edges on each time step also constrains the amount (and thus the cost) of communication between nodes based on a local dynamic criterion.

The model we suggest offers unique opportunities to study not only dynamics of the resulting nodal time series, but also the dynamics and changing topologies of active edges, as constrained by the brain’s structural network. We view the changing structural subgraph as an integral part of the ongoing dynamics of the model rather than intermediate steps toward a final, evolved network. We begin to analyze this novel, structural, edge time series, and we discuss its potential utility in the context of brain modeling.

## II. RESULTS

### A. Model Description

The innovation we propose limits broadcasting along edges by restricting nodes to communicate with only a selected subset of their structural neighbors on each time step. Nodes must select *m* neighbors to communicate with, and the edges to those neighbors are considered ‘active’ for that time step only. While there are a number of possible criteria by which nodes could choose neighbors, in this paper, we primarily focus on phase-synchrony: nodes select their *m* most phase-synchronized neighbors for communication. The degree of synchrony between every pair of nodes is given by the phase difference between the two nodes. On each time step, the localized, instantaneous order parameter, 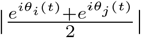, is calculated for every pair of connected nodes and stored in the matrix *R*(*t*). Each node then selects its *m* most synchronized neighbors, and the state of each edge (active or inactive) is stored in the binary matrix *δ*. The resulting network is symmetrized (so that edges remain undirected, see Methods and Discussion) such that *δ_ij_* = *δ_ji_* = max(*δ_ij_*, *δ_ji_*). The phase update is calculated according to the original Kuramoto-Sakaguchi equation. As such, the proposed model’s phase update is:

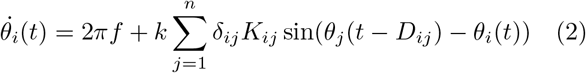

where *δ_ij_* is given by:

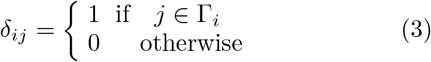

and Γ_*i*_ is the set containing the *m* most synchronised neighbors of node *i*, obtained by sorting the values in the i-th row of the matrix *R*(*t*). We note that Γ_*i*_ comprises only nodes for which *K* > 0, and is such that |Γ_*i*_| < *m* if *m* is larger than the degree of node *i*. Figure 1 compares one time step of the K-S model with one time step of the proposed model.

**FIG. 1.**
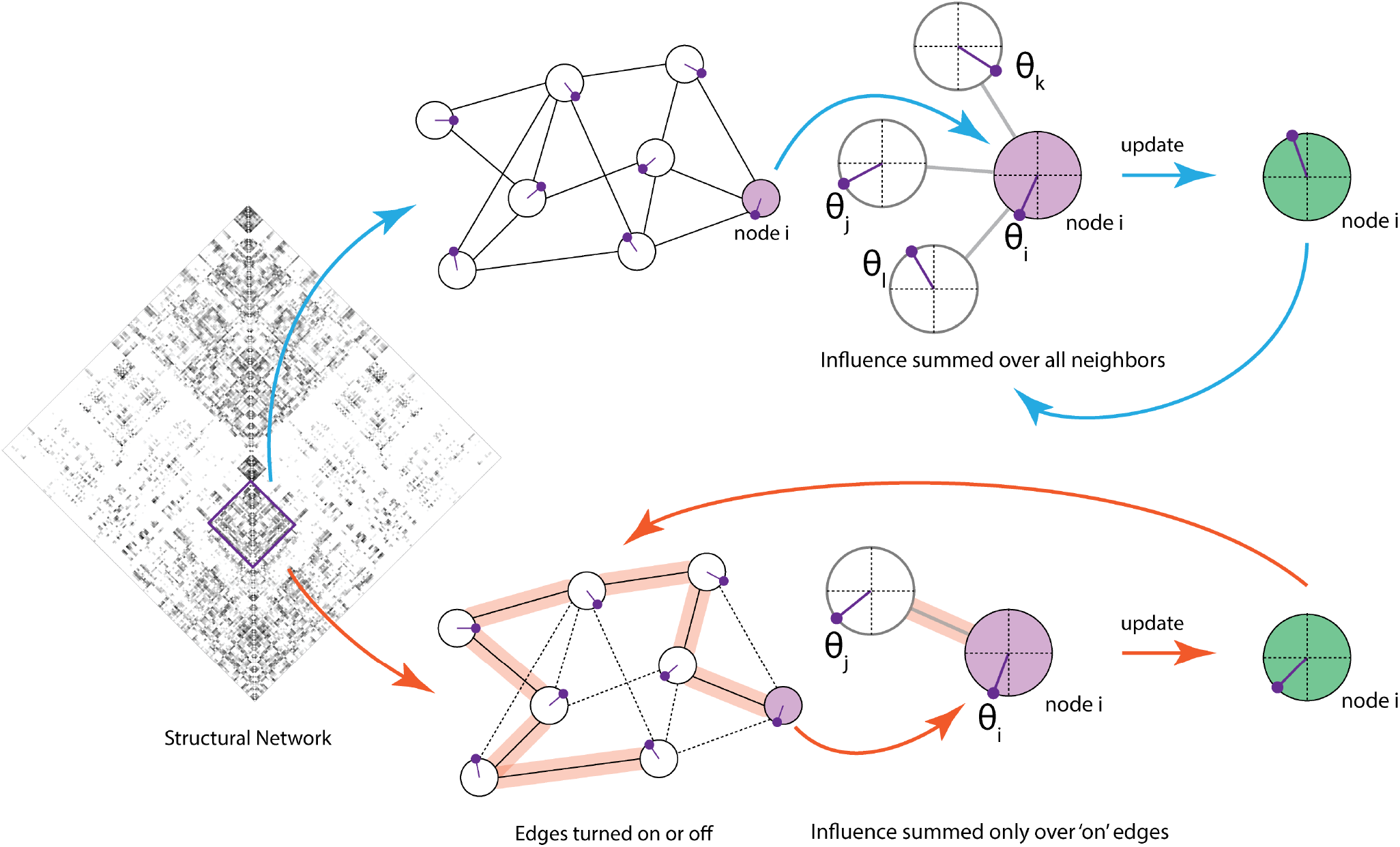
Schematic of single update of Kuramoto-Sakaguchi and the proposed model. The schematic shows the calculation of the phase update for a single node on a single time step of both the classic Kuramoto-Sakaguchi model (blue arrows) and the proposed variation of the model (orange arrows) for comparison. In the classic model, the update to node *i* is summed across all structurally connected neighbors according to 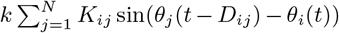, where *K_ij_* is the weight of the edge between node *i* and node *j*. In the proposed model, each node selects *m* neighbors with the smallest phase difference (most synchronized), and the corresponding ‘active edges’ are retained - all other edges are set to zero. Influence is then summed over the active edges, according to 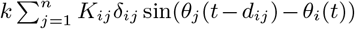, where *δ* is a binary matrix that stores whether edges are on or off.

Because the amount of phase-synchrony between neighbors is determined locally, this strategy does not require global information about the topology to guide edge selection, while imposing a more parsimonious and energetically-efficient communication strategy on the model. Note that turning connections on and off in this way restricts the selected structural network to be a subgraph of the full network. As *m* increases, this subset approaches the full, static network. It is also important to note that nodes are *required* to choose the full *m* number of neighbors. Even neighbors that have a large phase difference must be chosen if there are no other more synchronized neighbors.

Implementing the model in this way creates a timevarying network structure with a two-way relationship between model dynamics and network topology. The network chosen on each time step is a direct result of the phase of the oscillators on that time step and directly influences the phase update for the next time step. Creating this loop requires one additional parameter, *m*, but also outputs an additional, structural edge time series [32] of active edges for each time step. Next we explore how different settings of *m* influence the fit of modeled functional connectivity with empirical data.

### B. Empirical Fit

The proposed model allows structure and function to co-evolve according to two parameters, *k,* the global coupling constant, and *m*, the number of neighbors per node. We explored this parameter space by performing 13 runs for each set of parameters within the ranges 1 < *m* < 113(the maximum degree of the network) and 100 < *k* < 500. The range of coupling values was chosen in accord with previous work [31]. Because the model’s dynamics are chaotic, and the initial conditions as well as the intrinsic frequency of each node are chosen randomly, multiple runs are needed for each set of parameters to ensure that the results are not dependent on initial conditions. The model used a consensus-averaged anatomical network derived from 95 subjects of the Human Connectome Project (HCP) [34] as its full structural network (see Methods). Only edges that exist in this network could be turned on or off on a given time step. The resulting oscillator phase time series were convolved with a haemodynamic response function and down-sampled to match the time resolution of the resting state BOLD fMRI scans from the same subjects. Functional connectivity was calculated from the simulated time series and then correlated to the subject-averaged empirical functional connectivity to assess model fit. Results obtained by varying the parameters *k* and *m* are shown in Figure 2A. For many combinations of the two parameters, the empirical fit is significantly larger than what is obtained with the traditional K-S model’s single parameter, *k*. The proposed model resolves to the K-S model when *m* ≥ the maximum degree of the network. Here, the model becomes equivalent to the K-S model at *m* = 113, and so the parameter space of the traditional model (which varies only along *k*) can be seen in the last column of Figure 2A. In keeping with previous work [31], we focus on *k* = 280 in the subsequent analyses of this paper. The variation in performance along *m* with *k* = 280 is shown in Figure 2B in more detail.

**FIG. 2.**
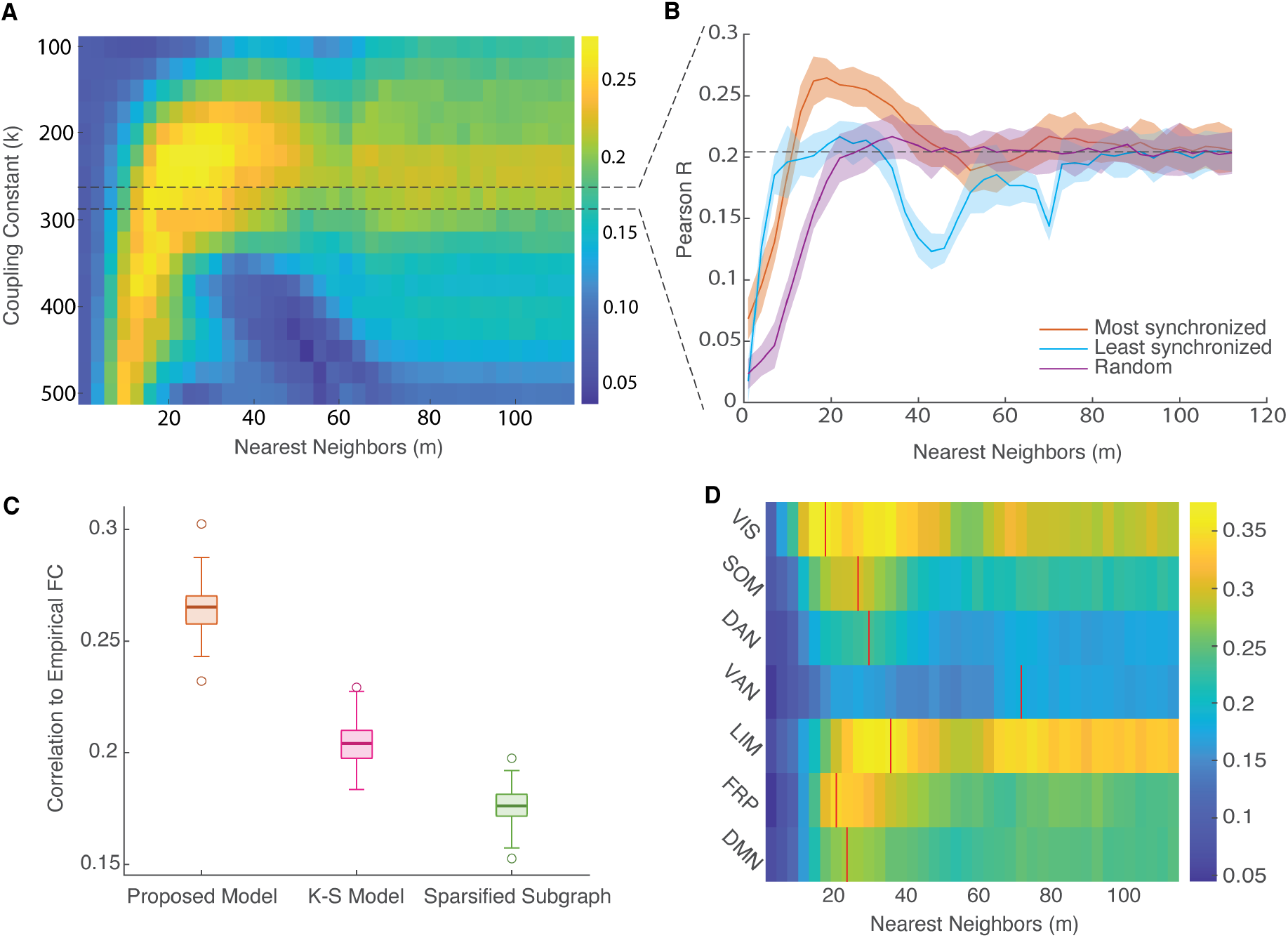
The proposed model shows improved empirical fit over a rich parameter space. A. The parameter space of empirical fits of the proposed model, using most synchronized neighbors as the edge selection paradigm. The number of neighbors selected per node, *m,* is shown on the x-axis, and the coupling constant *k* on the y-axis. Colors represent Pearson’s R with empirical FC averaged over 13 runs. Coupling constant *k* = 280 (used in ref [31]) was chosen for subsequent analysis, and is displayed with varying *m* values in panel **B**. **B** also shows two other edge selection paradigms, run with *k* = 280. The selection of *m* least synchronized neighbors is shown in blue, and *m* randomly chosen neighbors in purple. Clouds show a 95% confidence interval, and dashed line shows the mean performance of the Kuramoto-Sakaguchi (K-S) model over 200 runs. **C.** Boxplots comparing the model fits for 200 runs each of the proposed model with most synchronized edge selection (orange), the K-S model (pink), and the K-S model run on an SC matrix sparsified by retaining the strongest edges (green), with the density set to correspond to the average network density throughout a run of the proposed model. **D.** Modeled FC correlated to empirical FC for each of seven canonical functional systems [33] separately. X-axis shows variation along the number of nearest neighbors, and color represents correlation to empirical data. Each system’s peak correlation to its empirical FC is marked by a red line. All correlations computed as Pearson’s R.

With *k* = 280, the model reaches peak performance when *m* = 19, significantly outperforming the K-S model (two independent sample t-test, 398 degrees of freedom: *t* = 67.03, *p* = 6.4 × 10^−219^, Figure 2B-C). We also tested whether this model trivially improves empirical fit by expressing the underlying structural connectivity (which is significantly correlated to empirical FC), but this does not explain the improved fit (see SI figure 1).

To examine the effect of the edge selection strategy on model fit, two other selection criteria were tested for all m. In one alternative strategy, nodes selected their *m* least synchronized neighbors, and in the other, nodes selected *m* neighbors randomly. Neither strategy matched the performance of the model in which nodes selected their most synchronized neighbors (Figure 2B), indicating that the choice of edge selection strategy is relevant to empirical fit. However, both strategies reached their own best performance at *m* much smaller than the maximum degree of the network. Selecting the least synchronized neighbors matched the performance of the K-S model at *m* = 10, but performance fluctuated for larger *m* values. Randomly selecting neighbors matched the performance of the K-S model at *m* = 20, and performance did not worsen for larger values of m. This indicates that, even when edges are chosen at random, very few edges are needed on each time step to model empirical FC.

By selecting edges at every time step, the proposed model has the capability to greatly vary the active subgraph throughout a run, but is not forced to do so. It is possible that the improvement in empirical fit could be due only to the sparsification of the structural network, and not the exploration of the space of possible dynamic subgraphs of active edges. To test this, we compared the peak performance of our model to the performance of the K-S model run on a static, sparsified structural network. The sparsified network was created by re-weighting the SC network by how much time on average each edge was active in the proposed model. This network was then thresholded to match the average network density of the selected subgraphs in the proposed model. A total of 200 runs were performed for each of these three models, and boxplots of the correlation of modeled FC to empirical FC are shown for each model in Figure 2C. The proposed model significantly outperforms the K-S with a sparsified subgraph (two independent sample t-test, 398 degrees of freedom: *t* = 103, *p* = 4.94 × 10^−289^).

Finally, we asked whether a few canonical functional systems [33] may be driving the improvement in empirical fit. Correlation to empirical FC broken down by the seven functional Yeo systems is shown in Figure 2D. Six out of seven functional systems either display a peak or reach a plateau of performance at *m* ≈20. This suggests that the improved fit for *m* = 20 is not limited to a specific or very few functional systems, but is a network-wide phenomenon.

The Kuramoto Order Parameter 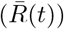 is an instantaneous measure of the synchrony of the system [20]. The mean 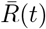 over time can be used to characterize the global synchrony of a system across an entire run. The standard deviation of 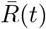 over time is thought to be an approximate measure of the metastability of a system [28], or the extent to which it explores its dynamical state space without settling into an attractor. It has been argued extensively in previous work that the brain is indeed metastable [29, 30, 35], and much work has shown metastable behavior in Kuramoto models as well [10, 28, 31]. We compared the mean 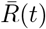 and standard deviation of 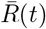 for all values of *m* at *k* = 280 for the three edge selection paradigms described above (Figure 3A,C). Least synchronized edge selection shows significantly greater global synchronization for many values of *m*, which is consistent with the literature [36]. However, when compared with Figure 2B, it becomes clear that choosing a model that maximizes global synchronization (mean R) or metastability (std R) does not necessarily increase empirical fit. Most synchronized edge selection shows relatively increased global synchrony for *m* = 20, where its best empirical fit was found, but it does not have the highest global synchrony found in this series of models. Finally, we compared the mean and standard deviation of 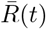 for 200 runs at the proposed model’s peak performance with the K-S model and the K-S model run on a static, sparsified subgraph, as before (Figure 3B,D). Both the mean and standard deviation of the order parameter appear to be situated between the values for the K-S model and the K-S model with the sparsified subgraph, indicating what may be a dynamic ‘sweet spot’ combining intermediate global synchrony with intermediate metastability.

**FIG. 3.**
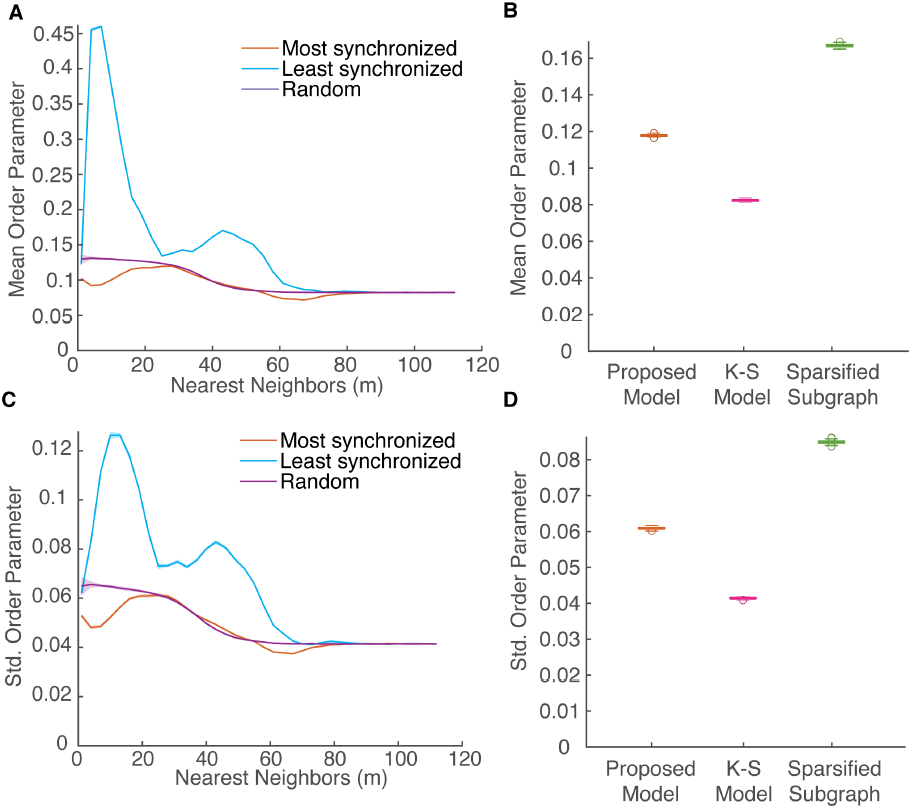
Synchronization and metastability vary with the number of nearest neighbors. A. The time-averaged Kuramoto Order Parameter 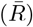 across *m* values for most synchronized, least synchronized and random edge selection paradigms. All runs performed with *k* of 280, and curves are shown with a 95% confidence interval. **B.** Boxplots showing the time-averaged order parameter for 200 runs each of the proposed model, the K-S model and the K-S model on a sparsified SC subgraph. **C.** The standard deviation of 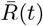 across runs with varying *m*. **D.** Boxplots showing standard deviation of 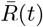 as in **B**.

### C. Structural Edge Time Series Analysis

In addition to the simulated BOLD time series, the proposed model also creates a novel structural edge time series of the active edges at each time point. In Figure 4, we summarize the activity of each edge in two ways. We count the number of times an edge switches state (active to inactive or vice versa), and divide it by its opportunities to flip (all time points in the run) to get a measure of how dynamic each edge is. We also calculate the number of time steps each edge spends active as a fraction of the total number of time steps in the run. These values were averaged over 13 runs of the model and are shown in node-by-node matrices as colored edge weights. The matrices are organized by structural modules, such that node pairs belonging to the same module are shown in blocks along the matrices’ main diagonals (see Methods). Ordering the nodes in this way reveals a static core of within-module edges that emerges for intermediate values of m. We further investigated the relationship between the structural modules and edge dynamics by showing the distribution of values for fraction of time active and flips for between and within module edges. For all values of *m* shown, edges within a module spend significantly more time active than edges between modules. For all shown values of *m* greater than one, between module edges flipped significantly more than edges within modules. Importantly, the scale of the flips matrices and violin plots should be noted: edges rarely change state, and the most dynamic edges change on only 1.5% of time steps, reflecting a network structure that evolves relatively slowly.

**FIG. 4.**
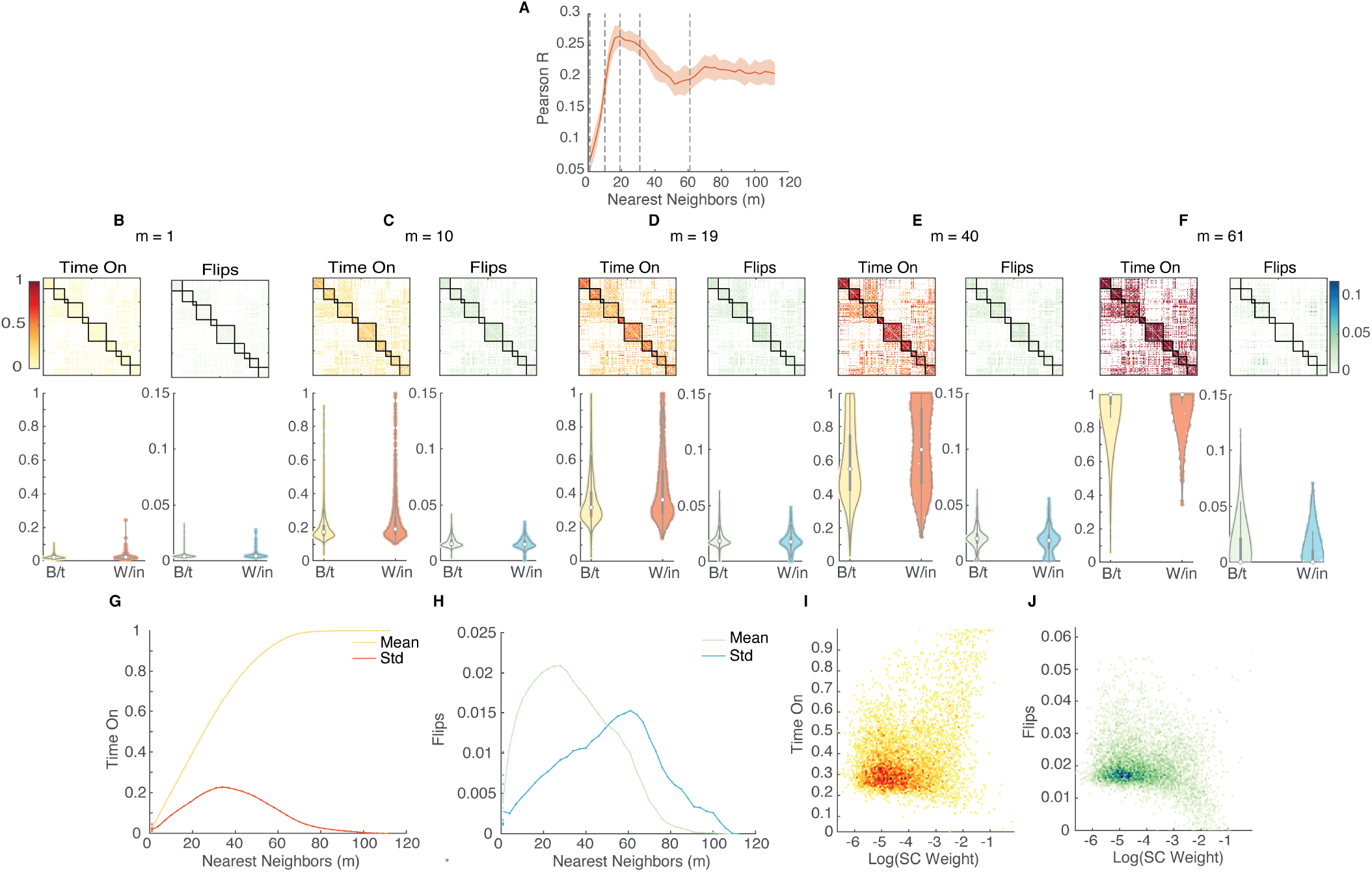
Variation in the amount of time an edge spends on and the number of time it flips is seen across *m*. **A.** Reproduction of Figure 2B. Dashed lines show the values of *m* at which panels **B - F** were observed. **B-F**. The amount of time each edge spent on and the number of times each edge flipped as a fraction of the total number of time steps is shown in node-by-node matrices where color represents the number of flips and amount of time on for each edge. Nodes are ordered by participation in structural modules (see Methods), which are marked along the diagonal. Violin plots show the distribution of number of flips and fraction of time on for each edge, separated by whether the edge lies between (B/t) or within (W/in) a structural modules. Figures show *m* values of 1, 10, 19, 40, and 61. At all values of *m* within-module edges spend significantly more time on than between-module edges according to the Mann-Whitney U-test (*m* = 1: *U* = 5.79; *m* = 10: *U* = 10.26; *m* = 19: *U* = 11.76; *m* = 40: *U* = 9.71; *m* = 60: *U* = 4.74, *allp* < 2.11 × 10^−6^). Between module edges also flipped significantly more than within module edges for all values of *m* except for *m* = 1, for which within module edges flipped significantly more (*m* = 1: *U* = 8.37; *m* = 10: *U* = −6.36; *m* = 19: *U* = −7.76; *m* = 40: *U* = −8.68; *m* = 60: *U* = −5.06, all *p* < 4.12 × 10^−7^). **G.** The mean and standard deviation of the amount of time on across all edges for every value of *m* at *k* = 280. **H.** The mean and standard deviation of the number of flips across all edges for every value of *m* at *k* = 280. **I-J**. The fraction of time on and flips is plotted against the log of the weight of the edge in the structural connectivity matrix. The correlations are significant, but weak (Time On: Spearman’s *ρ* = 0.14, p-value = 9.47 × 10^−27^, Flips: Spearman’s *ρ* = −0.16, p-value = 1.54 × 10^−34^), indicating that there are also topological effects at play.

While Figure 4B-F highlight in detail the edge structure for specific values of *m*, the mean and standard deviation of time active and flips across all edges for all values of *m* is also summarized in Figure 4G and H. As expected, mean time active increases monotonically with *m*, while standard deviation of time active peaks and then decreases. Consistent with results shown in Figure 2C demonstrating that the dynamic edge structure is necessary for improved empirical fit, only the mean number of flips has a peak coincident with the model’s peak performance at *m* = 20.

We tested whether the alignment of active edges to modules could be trivially explained by the edge’s weight. Within-module edges tend to be stronger than between-module edges, and therefore may be easier to synchronize, leading them to be chosen as active edges on more time steps. We plot the fraction of flips and time active against the log of the structural edge weight in Figure 4I-J. We find only a weak correlation (time active: Pearson’s *R* = 0.3, *p* = 4.4978 × 10^−126^, flips: *R* = −0.1897, *p* = 4.9134 × 10^−50^) between structural weight and time active or flips. This indicates that the relationship between modules and active edges is indeed topological, since much of this effect is not explained by edge weight alone.

So far, our analyses have explored the impact of varying the parameter *m* on the behavior of our model. Next, we focus on *m* = 19 to provide in-depth analyses of the model’s dynamics in its peak empirical performance. Due to computational and time constraints, each of these analyses were performed on only one run of the proposed model. While specific results may vary for other runs with different random initializations, the results shown here may be thought of as a first exploration into the model’s best performance.

Edges were clustered using a multiresolution clustering algorithm [37] (see Methods) to reveal two large edge communities that change (flip on/off) together in time (see Methods). The overall community structure appears non-assortative, instead displaying core-periphery organization (Figure 5A). A strong core community with similar time series can be seen at the top of the diagonal in Figure 5A. The other large community forms a peripheral structure defined by the edges’ relationship to the first community, rather than to each other. The edges belonging to each community are shown in Figure 5 in node-by-node matrices, where the edge between nodes *i* and *j* appears colored if it belongs to the community. These matrices are ordered by the structural modules, where nodes belonging to the same modules are placed together (see Methods). Membership in the first community is largely reserved for intramodular edges, with a high average time active (Figure 5B), consistent with Figure 4. The next communities are more diffuse, showing progressively lower average time active across the community.

**FIG. 5.**
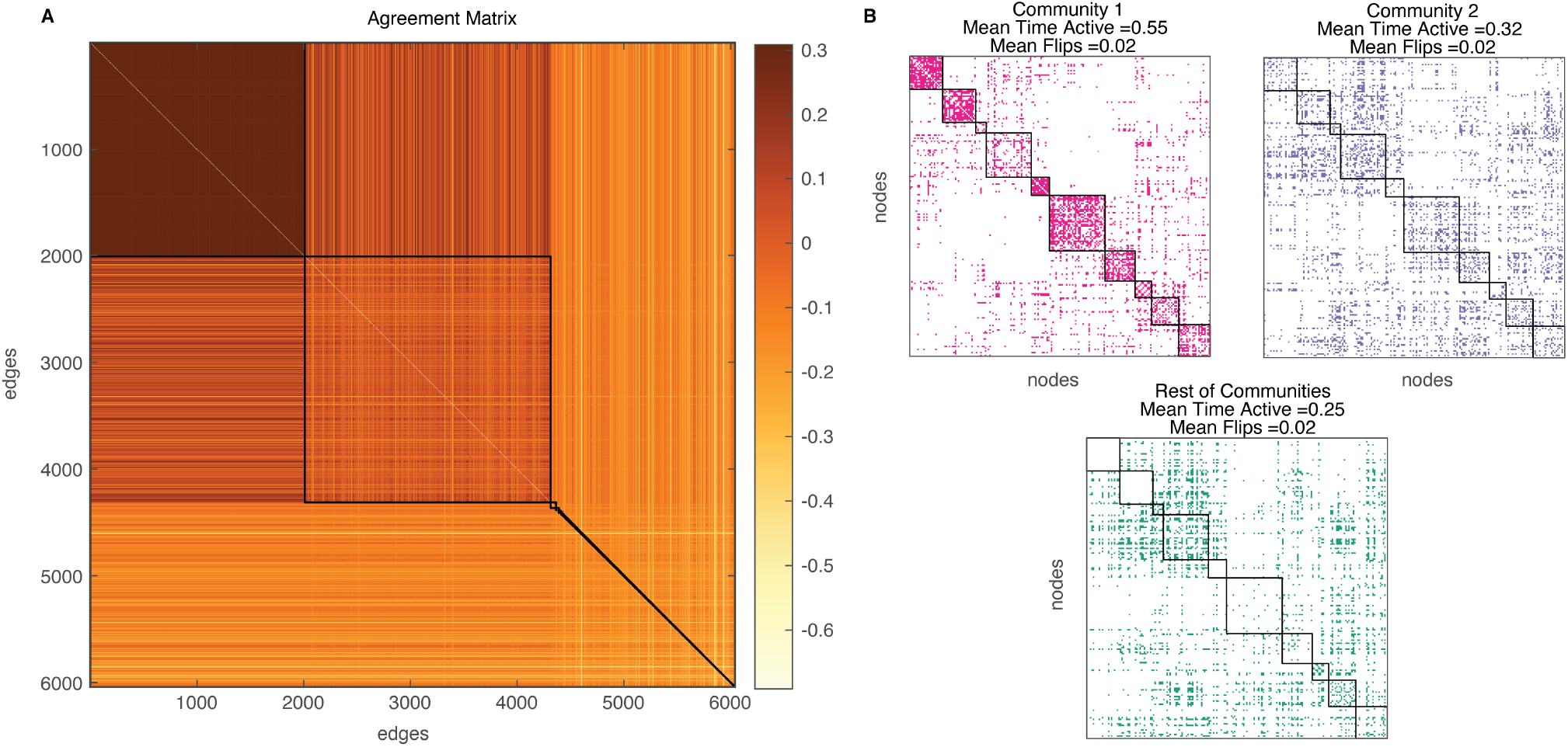
Edge clustering reveals core-periphery structure. The time series of all edges were clustered using multiresolution consensus clustering (see Methods). **A.** The edge-by-edge agreement matrix of 300 partitions sampled across resolution parameters. Two large communities are are found with a strong core-periphery structure. **B.** Communities found by the edge clustering are marked by different colors on node-by-node matrices. The edge between nodes *i* and *j* appears colored if it belongs to the community listed above the matrix and white otherwise. The matrices are organized by participation in structural modules shown on the diagonal (see Methods). The mean time active and mean number of flips for each edge community is also shown, revealing that the amount of time edges spend active consistently decreases for each community.

Finally, a suite of network metrics were calculated on the active subgraph at every time step to track the evolution of its topology. We found that all network metrics evolved much more slowly than the integration time step, with autocorrelation functions that decayed to zero at around 45 time steps (Figure 6A). The same slow decay is shown in a recurrence analysis, in which the mutual information between the binary subgraph on each point is calculated (see SI figure 2). The slow variation in network topology is dependent on the choice of edge selection strategy. The autocorrelation of the same set of network metrics calculated on a time series in which edges were chosen randomly decays to zero almost immediately (Figure 6A). An example of slow evolution can be seen in a small sample of the modularity time series (Figure 6B), which shows notable excursions away from the median of the time series. We further examined these excursions by plotting the complementary cumulative distribution function (CCDF) of the length of time of each excursion in Figure 6C, showing a heavy tail of excursion lengths. This CCDF is shown alongside a CCDF of the same time series that has been shuffled in time to demonstrate that the temporal structure is key to observing the heavy tail. To investigate the dependence of this result on edge selection strategy, the same CCDF was calculated for the modularity time series of a run in which edges are chosen randomly. The behavior of random selection is very similar to the temporally shuffled data, indicating the dependence of the heavy tail on edge selection strrategy. Taken together, these analyses indicate that choosing the most synchronized edges on every time step enables our model to evolve through distinct periods of low modularity (integration) and high modularity (segregation), as structural modules are created and dissolved in co-evolution with the oscillatory dynamics of the model.

**FIG. 6.**
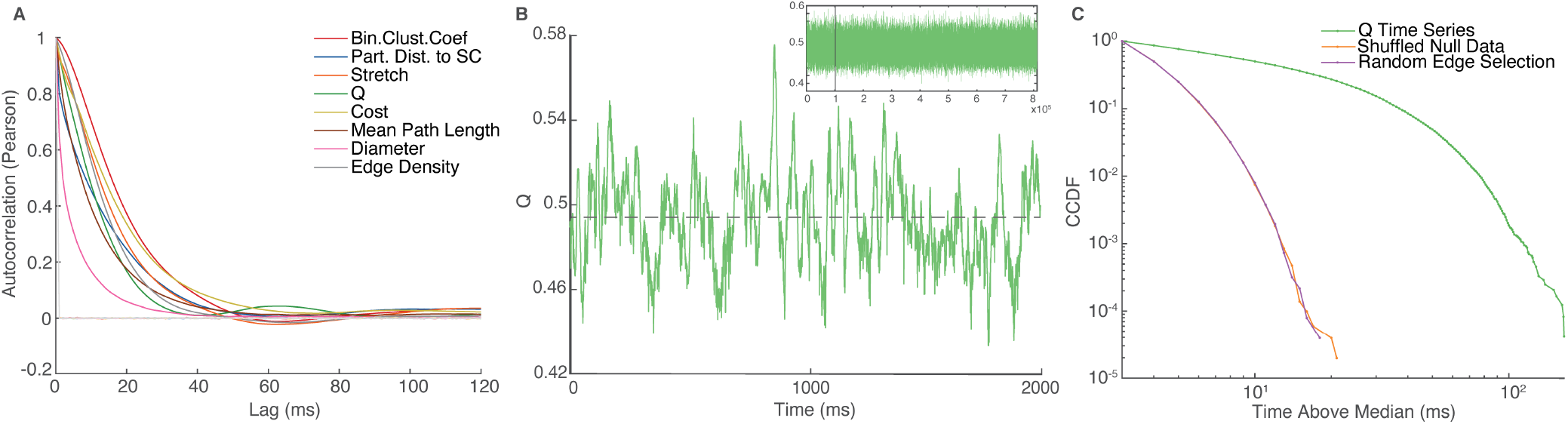
Network metrics of active subgraph evolve slowly over time. A. A suite of network metrics (see Methods) was computed on every time point in the structural edge time series of two runs of the model. One run employed most synchronized edge selection and the other employed random edge selection. The autocorrelation is plotted for each of these time series. The legend shows the name of the network metric. The autocorrelation of network metrics for the run selecting most synchronized edges is plotted in the color shown on the legend, while the same metrics for the run employing random edge selection are plotted in the same, but less saturated color. Random edge selection does not show long autocorrelation times. **B.** A small sample taken from the full time series (inset) of the modularity of the active subgraph (Q). The time series shows long excursions above and below the median (marked with a dashed line). **C.** Complementary cumulative distribution function of the length of periods of time that Q spends above its median value. Three time series are shown: Q time series from choosing the most synchronized edges, that same time series shuffled in time, and the Q time series from choosing random edges on each time step. A strong heavy tail is only seen when edges are chosen by selecting the most synchronized neighbors.

## III. DISCUSSION

### A. Contribution and Relationship to Previous Work

Here we implemented and explored the properties of a model that limits communication between nodes on each time step and allows its structural topology and dynamic state to co-evolve through time. Functional connectivity derived from the model achieves significantly greater fit to empirical data, when compared to the standard K-S model. Furthermore, a novel analysis of the evolving graph of ‘active edges’ further highlights the importance of structural modules and reveals the emergence of slowly evolving structural network topology, each topology representing a different configuration of signaling pathways between nodes. As such, the model provides an initial link between studies focusing on dynamic neural mass modeling and network communication.

The improvement in empirical fit seen in the proposed model cannot be achieved without a dynamic edge structure and appropriate rules governing its evolution. Evidence for the importance of a dynamic edge structure is seen in Figure 2C, as the K-S model run on a sparsified, static subgraph could not match the performance of the K-S model, much less the performance of the proposed model. Randomly choosing edges creates a network with the same edge density as the sparsified subgraph, but varies that network on every time step. The additional structural edge dynamic improves performance to match that of the K-S model. In order to improve performance above that of the K-S model, not only dynamic edges, but a good rule determining their selection must be incorporated. As seen in Figures 2B and 3, changing the edge selection strategy changes the model’s dynamics and performance significantly. No other edge selection rule could match the peak performance achieved by choosing each node’s most synchronized neighbors. The random selection of edges on each time step can be seen as a null model for *any* edge selection rule, as it creates a dynamic structrual subgraph without order. Randomization cannot match the performance of choosing each node’s most synchronized neighbors. However, it does match the performance of the K-S model long before all edges have been incorporated (Figure 2B), and even before *m* approaches the average degree of the network (63 edges). This provides strong evidence that not all edges are created equal: having more active edges in the model does not directly translate to better model performance. Our work suggests that the exploration of new model mechanisms can lead to improved ability to predict function from structure.

The proper choice of edge selection rule is also essential for the emergence of order in the time-varying structure of the model. When edges were chosen randomly, emergence of slowly evolving network metrics or periods of high and low modularity was not observed. These transient features arise organically out of local edge selection rules and are not necessarily expected from the model’s design or parameters. Self-organizing topological features from local plasticity rules have been described in other studies [38], and the phenomenon of transient emergent structure has been explored in information-theoretic contexts [39]. This study highlights the importance of the underlying mechanism of edge selection to emergent dynamics and creates a framework for computational explorations of alternative strategies and model formulations.

The mechanism of edges switching between active and inactive states, in addition to coupling structure and function, allows the model to perform with a higher energetic efficiency than other dynamic models, which is a priority in communication modeling. While we do not implement energy cost explicitly in our model, we argue that our model’s operation is conceptually more efficient than other models of coupled differential equations. At a microscopic level, each action potential requires energy to depolarize and repolarize a neuron, a demand thought to ultimately shape energy requirements for inter-areal brain communication [13, 40–42]. Dynamic influence along edges, as implemented by the K-S model, should require a similar metabolic expenditure. By restricting the number of edges available to the system on every time step, we limit the number of channels capable of relaying dynamic influence, and thus reduce the resulting metabolic cost.

The model also makes few demands on the ‘informational cost’ required to implement its edge selection strategy. It has been argued that communication strategies that require global knowledge of the network topology are less biologically plausible than those requiring only knowledge of the local neighborhood [13]. The broadcasting strategy employed by the K-S model uses edges indiscriminately and requires no knowledge of global network topology. Our model adds only a modest additional cost by implementing edge selection using a rule based on local information about synchrony. Hence, while imposing higher informational cost than the standard K-S model, it is still solely dependent on local information that could be conceivably accessed by individual network nodes.

Of the local edge selection strategies explored here, selecting each node’s most synchronized neighbors is the most biologically plausible. As well as requiring little topological information, the rule is in accord with literature suggesting synchrony as a requirement for neural communication. The communication through coherence hypothesis holds that neurons communicate best when synchronized with each other [35, 43]. Our model explores many possible synchronization states, and so also explores a great number of possible structural subgraphs (see SI Figure 2). The combination of metastability (the exploration of many dynamical states) and edge selection echoes the concepts explored in ref [35] describing how the communication through coherence hypothesis may be implemented in dynamic models of neural activity. Additionally, empirical work has shown that the state of local oscillations affects the ability of neural populations to respond to a stimulus [44]. By relying on measures of local synchrony, our model may replicate this behavior: regions that are not sufficiently synchronized are unable to pass signals to each other.

Metastability and global synchrony have been important focal points of past modeling studies of brain dynamics [28–30, 35, 45, 46]. Our work suggests that simply maximizing metastability may not be sufficient for accurately reproducing features of empirical functional connectivity. The analysis presented in Figure 3 demonstrates that the version of the model with the greatest average metastability and global synchrony (the K-S model run on a limited subgraph) produced relatively poor empirical fit (Figure 2C). While the proposed model does exhibit metastability and global synchronization above that of the K-S model, and while these properties may contribute to its improved performance, the decreased empirical performance of the K-S model employing the static, sparsified, subgraph suggests the need for a more nuanced interpretation of the importance of these two criteria.

Dynamics of networks and the analysis of temporal networks has been of great interest across complex systems theory and multiple applied fields of study [47–51]. In many real-world networks, the network topology and link structure changes through time, in processes such as contagion, social exchanges, or neuronal signaling. Indeed, past computational work has sought to create models with co-evolving structure and dynamics [38, 52–58]. Variations on the Kuramoto model form a small subset of models coupling the evolution of structure and function. Some Kuramoto model variants have implemented Hebbian growth rules [59, 60], while others break and rewire connections over time [36, 61]. In studies using Hebbian rules, it was found that the resulting dynamics of the model can be altered significantly by implementing different variations on Hebbian learning, which parallels findings reported in our study. However, Hebbian plasticity allows only the weights of a network to change, while the topology remains constant, which entails that most of the graph-theoretic analyses carried out here are inapplicable. Additionally refs. [59] and [60] found that Kuramoto models implemented with a Hebbian learning rule settle into a steady state with unchanging weights. We did not observe similar behavior in our model, likely due to the combination of intermediate coupling, which allows the model to explore a larger number of synchronization states, and the sensitivity of the edge selection rule to small changes in the individual phases of the oscillators. However, for the purposes of modeling ongoing brain dynamics at rest, we view a model that is capable of generating continuous dynamics without being driven by external input as advantageous.

More recently, Papadopoulos et al. [36] proposed a variation on the Kuramoto model that is similar to what we introduce here. In deleting and stochastically rewiring the most synchronized edges at set time intervals, they find that the structural network employed in the model evolves topological properties, such as degree-frequency offset correlations, that support a greater level of global phase synchrony than the initial network. This accords with the high levels of global synchrony we found when nodes chose the least synchronized neighbors during edge selection. Our study differs from Papadopoulos et al. primarily in the goals and framework of the study. They view rewiring edges as a self-organizing method of evolution toward a final network topology, whereas here we have focused more on the ongoing dynamics of edges, situated specifically in a neuroscientific context. While the edge update rule that they introduce is different from any introduced in our study, their rule (and many variations on it) could also be implemented in our model and compared to empirical data. Its element of stochasticity may change the model’s behavior in surprising ways. Both papers demonstrate a need for more exploration and understanding of dynamics of networks and its effects on dynamics on networks, especially through computational modeling.

Our work is also complementary to recent interest in time-varying functional connectivity in neuroimaging studies. We hope that our model may add a new lens through which to explain dynamic phenomena observed in windowing approaches [62] and more recent edge-centric approaches [32]. A particular area of interest may be the temporal emergence of periods of high and low modularity found in our analysis of structural edge time series. Extensive empirical work has shown the existence of similar periods of segregation and integration in functional brain data [63–67], including work that demonstrates that functional modules can reconfigure across time [68, 69]. How such fluctuations observed in nodal time series may reflect underlying changes in effective or active connections is a topic for future investigation. One hint of a link between these domains of variability may come from our observation that fluctuations in ‘active edges’ occur in specific patterns. We found that within-module edges formed a static, unchanging core with a dynamic periphery of more variable inter-module edges. This echoes what has been found in previous empirical studies of neuronal dynamics showing that time-varying functional connectivity reveals a static, modular core and a dynamic periphery [70, 71]. It is important to note that large changes in topological properties may require only small changes in the subgraph, at times only a single edge. Many of these changes are missed by coarse-grained analyses, such as we have performed here. Much more detailed analyses of the structural edge time series are possible and the goal of future work. For example, more thorough analyses may seek to make connections between oscillator dynamics and edge switches, as well as examine information flow throughout the network, and how such flow is responsive to the changing network topology.

One of the greatest limitations of the proposed model is that while it restricts the diffusive dynamics of the K-S model in a principled and informationally efficient way, it does not change the fundamental communication process. The model continues to employ coupling terms over all active edges present in the subgraph. While this subgraph is limited and time-varying, the communication mechanism acting on top of it remains diffusive in nature. Perhaps by creatively designing a new edge selection procedure or importing the edge selection framework to a different model, future work will be able to accurately model the diverse set of communication strategies that this work was inspired by, such as routing by shortest path lengths[72], search information[14], or navigation [19]. While changing the edge selection paradigm is not perfectly analogous to implementing a different communication strategy, a key result of this work is that varying the edge selection rule changes model dynamics and performance significantly. This is promising for the future development of models truly capturing the moment-to-moment unfolding of various communication strategies. Our work represents an initial attempt to bridge the gap between two large classes of models of brain function, based on dynamics and communication.

### B. Technical Considerations

On every time step the chosen subgraph was symmetrized so that it remained undirected, as the original network was also undirected. We chose to implement this in such a way that an edge only had to be chosen by one of its incident nodes in order to be considered active on the time step. It was inconsequential for an edge to be chosen by both of its nodes. This resulted in a small variation of edge density across time that is reported in Figure 6A. Another consequence of this implementation is that node degree remains somewhat variable throughout the chosen subgraph, and that the parameter *m* actually represents a lower bound on the degree of each node.

We also chose to implement edge-switching as a discrete process rather than a continuous one. As discussed above, there are several versions of the Kuramoto model that implement a continuous Hebbian-like learning process on the edges of a network [59, 60]. However, neuro-biologically, Hebbian mechanisms occur at the level of individual synapses, not large anatomical fiber tracts, from which the networks used here are derived. By activating and inactivating edges discretely, we situate our model more closely to communication models, in which communication events are thought to happen as discrete signals, rather than continuous interaction. Additionally, there is support for the importance of discrete information routing mechanisms that arise from underlying continuous dynamics [73, 74]. Our model could be thought of as abstracting away the underlying continuous dynamics to model the discrete information switching described by ref [73] explicitly. Finally, there is an element of simplicity and parsimony in flipping edges discretely as opposed to introducing an equivalent number of equations to the dynamical system.

Another possible edge selection strategy not explored here would be to create a global threshold for *R*(*t*). Every edge synchronized above threshold would be active for that time step. We do not explore this edge selection strategy because limiting the number of neighbors (rather than the degree of synchronization), requires relatively desynchronized nodes to maintain contact with at least *m* other nodes, thus preventing the formation of small, detached components. In this scenario, a small, highly synchronized cluster of nodes could dominate the subgraph’s active edges, forcing desynchronized nodes out of the network and creating a feedback loop in which they become further desynchronized from the rest of the oscillators. Requiring the choice of *m* neighbors assures continuing phase heterogeneity in the system, essential for modeling ongoing brain dynamics.

### C. Conclusions

Here we have shown that limiting the number of edges available to a set of Kuramoto oscillators with dynamic local rules leads to a better simulation of empirical functional connectivity from structural connectivity. The mechanism of edge selection we employ here was inspired by the principles of communication models in its metabolic and informational efficiency, thus helping to create a link between two important approaches to modeling brain function. The improvement in empirical fit achieved by our model demonstrates dynamic modeling will benefit from further exploration of novel mechanisms and the coupling of structure and function. We have only begun to explore the dynamics of the structural edges produced by our model, and hope that future studies will take a deep dive into the way structure and function are able to co-evolve together.

## IV. METHODS

### A. Dataset

Structural and functional data were taken from 100 unrelated subjects of the Human Connectome Project (HCP). Detailed collection procedures can be found in [34]. All data were downloaded minimally preprocessed according to the methods described in ref [75]. All participants gave informed consent to study protocols and procedures approved by the Institutional Review Board at Washington University. A very brief description of data collection is as follows. A Siemens 3T Connectom Skyra furnished with a 32-channel head coil was used for data acquisition.

Participants underwent two diffusion MRI scans, collected with a spin-echo planar imaging sequence with a repetition time (TR) of 5520 ms, echo time (TE) of 89.5 ms, a flip angle of 78°, isotropic voxel resolution of 1.25mm, b-values of 1000,2000,3000 s/mm^2^, and 90 diffusion weighted volumes in each shell, 18b = 0 volumes. Each scan was taken with opposing phase encoding directions and averaged. Functional magnetic resonance imaging (fMRI) was acquired from four resting state sessions (duration 14:33 min, eyes open) performed on two separate days. Scans followed a gradient-echo echo-planar imaging (EPI) sequence with a TR of 720ms, a TE of 33.1ms, a flip angle of 52°, a 2mm isotropic voxel resolution, and a multiband factor of 8.

95 participants were selected for inclusion in the present study based on the following exclusion criteria determined beforehand. The mean and mean absolute deviation of the relative root mean square (RMS) motion were calculated across either one diffusion MRI scan or the four resting-state MRI scans, producing four motion measures. Subjects were excluded if they surpassed 1.5 times the interquartile range of the measurement distribution in two or more measures. These criteria resulted in the exclusion of four subjects. An additional subject was excluded due to a software error during processing of diffusion MRI. Subjects that were included were 56% female, with an age range of 22-36 and mean age 29.29 ± 3.66.

### B. Structural Preprocessing

Structural data from diffusion imaging were minimally preprocessed before they were downloaded [75]. Additionally, nonuniformity in the images was corrected for using N4BiasFieldCorrection [76]. FSL’s dtifit was used to acquire mean diffusivity, mean kurtosis and fractional anisotropy scalar maps. Using these maps in conjunction with the Dipy toolbox (version 1.1) [77], a multishell, multitissue constrained spherical deconvolution [78] with a spherical harmonics order of 8 was fit to the diffusion data, using tissue maps estimated with FSL [79]. Probabilistic tractography [80] was performed nine times per subject using Dipy’s Local Tracking module. On each run, the step size (0.25mm, 0.4mm, 0.5mm, 0.6mm, and 0.75mm, with turning angle at 20°) and maximum turning angle (10°, 16°, 24°, and 30°, with step size at 0.5 mm) of the algorithm were varied. For each instance of tractography, each voxel of a white matter mask was randomly seeded three times with streamlines. In accord with Dipy’s implementation of anatomically constrained tractography [81], only streamlines longer than 10mm and with valid endpoints were kept. Further, errant streamlines were filtered out based on the cluster confidence index [82].

To derive subject-level structural connectivity networks, for each tractography instance the number of streamlines between each node of the Schaefer 200-node parcellation [83], in each subject’s anatomical space were recorded. Counts were then normalized by the geometric average volume of each node. A weighted mean based on the proportion of total streamlines at a given edge was used to aggregate edge weights across tractography instances for each subject. Performing the weighted mean in this way biases edge weights toward larger values, which reflects the tractography runs that were better parameterized to estimate the geometry of each connection.

From the structural connectivity network for each subject, a consensus SC network was derived for the entire group by doing a consensus average while conserving the global connection density to match the subject-averaged connection density, and maintaining the distribution of fiber lengths [84]. The network that resulted (200 nodes, 6040 edges, 30.4% density, 72.4 mm connection length) was used as the connection matrix on all simulations. This matrix was also used in ref [31], which provides more details on its construction. In the proposed model, edges were selected from this network to form subgraphs on each time point.

### C. Functional Preprocessing

Resting state fMRI data were minimally preprocessed before downloading [75], resulting in an ICA+FIX time series adhering to the CIFTI grayordinate coordinate system. Two additional preprocessing steps were performed: global signal regression and band pass filtering with a band from 0.008 to 0.08 Hz [85]. After confound regression and filtering, the first and last 50 TRs were discarded from the data, leading to a final scan length of 13.2 min, or 1,100 TRs. These surface-level functional data were averaged within each region of the 200-node Schaefer parcellation in CIFTI space [83] at each TR, forming 200 time series per subject.

To calculate subject-specific functional connectivity (FC) matrices, the 200 nodal time series were first z-scored. The Pearson correlation of each pair of nodal time series was calculated to produce a fully weighted, fully signed adjacency matrix. We averaged this matrix across all 95 subjects and then applied the fisher transformation to result in a single 200-by-200 empirical FC used for assessing empirical fit.

### D. Structural Modules

The structural connectivity matrix and the modules derived from it are identical to those used in ref [31]. These modules were detected using multiresolution consensus clustering, which captures communities across multiple scales [37, 86]. The Louvain method [87] was used for modularity maximization, with spatial resolution parameterized by *γ*. A parameter sweep across *γ* was performed twice. In the first sweep, with 1,000 steps, outer bounds yielding between two and *N* communities were established. A second sweep, with 10,000 steps over the same range, collected partitions found in the first sweep. Partitions were aggregated into a co-classification matrix, and a null model was subtracted from this matrix. The null model captured the expected degree of co-classification of two nodes based on the size and number of modules found. In the final step, the resulting matrix was clustered again using a variable consensus threshold *τ*. The results of different values of *τ* were examined, and the most frequently sampled partition was selected. This partition provides the ten structural modules examined in this study.

### E. Model Implementations

The results shown in this study are from two different implementations of the Kuramoto model: the standard Kuramoto-Sakaguchi (K-S) model and the proposed model.

The phase update of the K-S model is:

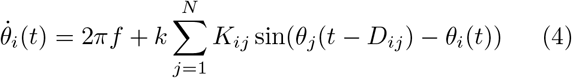

where *K* is a matrix that stores the weight of the structural connection between nodes *i* and *j*, *D_ij_* is the phase delay between nodes *i* and *j*, and *k* is a global coupling constant.

The innovation of the proposed model is to include a binary matrix, *δ,* re-calculated on every time step according to the edge selection rule, that activates or deactivates edges. The inclusion of *δ* changes the phase update as follows:

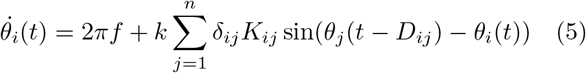

There are multiple ways to select *δ*, as we discuss in the following section.

The set of coupled equations describing the proposed model were integrated at 1 ms time resolution using a fourth order Runge-Kutta-Fehlberg method [88, 89] implemented in MATLAB. The mean intrinsic frequency of the oscillators was set to 40 Hz and varied among nodes according to a Gaussian distribution with standard deviation of 0.1 Hz. Time delays are implemented as phase delays, with a phase delay matrix computed from the physical length of each edge in the structural network, taken from a group average of the length of each edge’s constituent streamlines. The edge length is converted into a phase delay using a constant velocity. In this study, *v* = 12m/s is used, consistent with prior work [31]. This velocity leads to a mean time delay of 6.1 ms over all structural connections, however, it should be noted that the mean delay of the subgraph chosen at every time step may vary. The mean expected phase difference calculated using the intrinsic frequency is used to convert time delays into phase delays.

Initial phase conditions were chosen uniformly from the interval [0, 2*π*]. The initial structural subgraph was selected according to the edge selection rule on the initial phases.

Simulations were run for 812s of simulated time, allowing for a 20s transient to be discarded to match empirical scan time of 792s. The resulting phase time series was converted to amplitude by taking the *sin*(*θ*(*t*)), and then convolved with a canonical HRF function to produce a BOLD time series. The time series were low-pass filtered with a 0.25 Hz cutoff and then downsampled, averaging over 720 ms intervals of time, to correspond to the TR in the empirical rs-fMRI data. The resulting time series consisted of T = 1,100 frames, and underwent global signal regression to match empirical data. Residuals from the regression were kept and used to calculate simulated functional connectivity.

### F. Edge Selection Strategies

In this study we implement three strategies of choosing the matrix *δ*, which is incorporated into the K-S equation, and stores whether an edge is active or inactive.

For most of this paper we focus on the strategy in which nodes choose their *m* most synchronized neighbors. We calculate the order parameter, *R_ij_*(*t*) for every pair of oscillators *i* and *j* (see section G) to measure their phase synchrony. The entry of δ for an oscillator pair is then defined as:

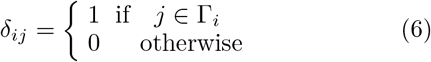

where Γ_*i*_ is the set containing the *m* most synchronised neighbors of node *i*, obtained by sorting the values in the i-th row of the matrix *R*(*t*). Γ_*i*_ can only contain nodes j for which a structural connection exists, i.e. *K_ij_* > 0. It is also such that |Γ_*i*_| < *m* if *m* is larger than the degree of node *i*.

We also compared the case in which nodes choose their *m* least synchronized neighbors for communication. Again, the order parameter, *R_ij_*(*t*) for every pair of oscillators *i* and *j* (see section G) is calculated to measure their phase synchrony. The matrix *δ* remains as defined above, except that Γ_*i*_ is now the set containing the *m* least synchronised neighbors of node *i*, as obtained by sorting the entries of row *i* of *R*(*t*) in descending order. Edges are again restricted to be present in the matrix *K*, and the cardinality of Γ_*i*_ cannot exceed the degree of node *i*.

Finally, we implement a strategy in which each node selects *m* neighbors randomly. In this strategy Γ_*i*_ contains a random selection of nodes *j* for which *K_ij_* > 0.

After edge selection is performed, in all three strategies, the resulting δ matrix is symmetrized to maintain undirectedness. As long as an edge is chosen by one of its incident nodes, it is retained. Edges chosen by both nodes are collapsed to form a single edge, so that *δ_ij_* = *δ_ij_* = max(*δ_ij_*, *δ_ji_*).

### G. Kuramoto Order Parameter

The Kuramoto Order Parameter is a measure of the instantaneous phase-synchrony of a set of oscillators, bounded between 0, when the oscillators are completely incoherent and 1 when the system is completely phase-synchronized. It is defined for any set of two or more oscillators. We calculate the instantaneous order parameter for every pair of oscillators during edge selection as:

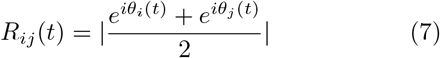

where *θ_j_* is the phase of oscillator *j*, and *i* is the imaginary number.

This value can be averaged over all oscillator pairs to obtain an instantaneous measure of the global synchrony of the system, given by:

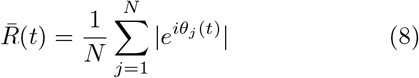

where *N* is the number of oscillators. Finally, this value can also be averaged over all time steps to obtain 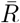, the average global synchronization of an entire run, as reported in Figure 3.

### H. Edge Clustering

Edges were clustered using multiresolution consensus clustering [37]. First the Jaccard distance between the time series of every set of edges was computed to create a 6040 × 6040 matrix. Louvain community detection was run 300 times on this matrix over a gamma range of 0-0.33. A co-classification matrix was formed from the resulting partitions, and a null model that accounts for the expected degree of co-classification between two nodes was subtracted. Finally, the co-classification matrix underwent consensus clustering with *τ* = 0 [86].

### I. Network Metrics

All network metrics were calculated using the Brain Connectivity Toolbox [90], which can be found at: https://sites.google.com/site/bctnet/. Helpful in-depth descriptions and explanations of these metrics can also be found in ref [18]. All metrics except cost were computed on the binary *δ* matrix for each time step.

#### 1. Binary Clustering Coefficient

The clustering coefficient is the fraction of a node’s neighbors that are also neighbors of each other, or the fraction of complete triangles around a node. We average this value across the binary (‘on/off’) subgraph at each time step:

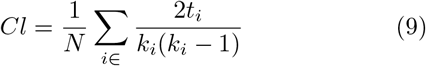

where *N* is the number of nodes in the network, *k_i_* is the degree of node *i* and *t_i_* is the number of closed triangles attached to node *i*.

#### 2. Shortest Path Length, Diameter, and Stretch

The shortest path between two nodes in a binary network is the path between two nodes that requires traversing the fewest edges. The BCT calculates weighted shortest path lengths using the Floyd-Warshall algorithm. Here, we calculate the mean of all shortest path lengths in the subgraph:

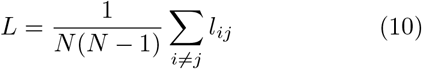

where *l_ij_* is the shortest path between nodes *i* ad *j*. The diameter is simply the longest shortest path.

The stretch quantifies the extent to which traveling between two nodes by diffusion requires more hops than traversing along the shortest paths, i.e. the mean first passage time. Put another way, the stretch quantifies how suited a network is to a diffusive process. Here we quantify “diffusion” as the mean first passage time between two nodes. We average over all node pairs and divide this value by the average shortest path length for the subgraph to arrive at the stretch. Formally,

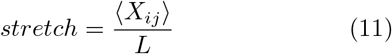

where 〈*X_ij_*〉 is the average mean first passage time between all pairs of nodes, *i* and *j*.

#### 2. Cost

The cost of the chosen subgraph is a measure of how expensive the edges in the subgraph would be to biologically form and maintain. It is given by the sum element-wise product between the weight and length of every edge:

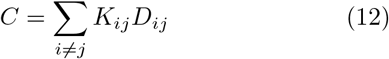

where *K_ij_* is the a matrix storing the weight of all edges and *D_ij_* is a matrix storing the length in mm of every edge.

#### 4. Modularity (Q) and Partition Distance to Structural Modules

Modularity (Q) is a statistic that quantifies the degree to which a network can be divided into non-overlapping communities that maximize the number of edges within each community and minimize the number of edges between communities. We used the Louvain algorithm [87] to find the optimal community structure and its associated modularity score (Q):

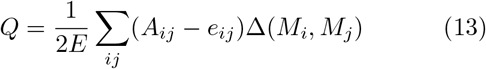

where *E* is the number of edges in the subgraph, *A_ij_* is the adjacency matrix of the subgraph, *e_ij_* is the number of edges expected by chance, given by 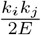, where *k_i_* is the degree of node *i*, and Δ(*M_i_*, *M_j_*) is the Kronecker delta function, which is equal to 1 if the nodes *i* and *j* are found in the same module, *M*, and 0 otherwise. The modularity was calculated for the active subgraph on every time step to create a modularity time series, which was used in further analysis (see section J).

The community partition found by the Louvain method can be compared to the structural modules found above by calculating the partition distance (the normalized variation of information between the two partitions, *X* and *Y*):

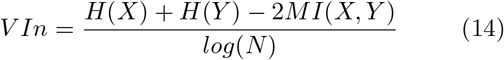

where *H* is the entropy of either partition, *MI* is the mutual information between both partitions, and *N* is the number of nodes in the network.

#### J. CCDF of the Modularity Time Series

To assess how the integration/segregation balance changed over time, we examined the distribution of long and short periods of high modularity across the run. The time series of instantaneous modularities was calculated as described in section I for the structural edge time series generated by selecting the most synchronized edges as well as the structural edge time series generated by selecting edges randomly. Both time series were binarized about the median and an excursion was defined as a temporally contiguous sequence of above-median bins (in a manner inspired by prior work on avalanche duration dynamics in neural circuits [91]). We computed the distribution of excursion lengths (i.e. how many excursions of length three, how many excursions of length five, etc).

We used complementary cumulative distributions (CCDFs) [92] to visualize the distribution of excursion lengths for both time series. CCDFs were chosen rather than histograms since they represent every data point, and decrease monotonically, rather than being visually disrupted by noise in the tails (for further discussion, see [92]).

To ensure that the distribution had a heavier tail than would be expected by chance, we compared the empirical CCDF to a permutation null, created by shuffling the modularity series in time.

## V. ACKNOWLEDGMENTS

M.P. is supported by the upported by the National Science Foundation Graduate Research Fellowship under Grant No. 2240777. Any opinion, findings, and conclusions or recommendations expressed in this material are those of the authors(s) and do not necessarily reflect the views of the National Science Foundation. M.P. and T.F.V. are supported by the NSF-NRT grant 1735095, Interdisciplinary Training in Complex Networks and Systems. The funders had no role in study design, data collection and analysis, decision to publish, or preparation of the manuscript. Data were provided, in part, by the Human Connectome Project, WU-Minn Con sortium (principal investigators: D. Van Essen and K. Ugurbil; 1U54MH091657), funded by the 16 NIH Institutes and centers that support the NIH Blueprint for Neuroscience Research and by the McDonnell Center for Systems Neuroscience at Washington University. M.P. would additionally like to thank her husband for countless warm dinners and hours of support.

