## Supplemental Information for "Co-Evolving Dynamics and Topology in a Coupled Oscillator Model of Resting Brain Function"

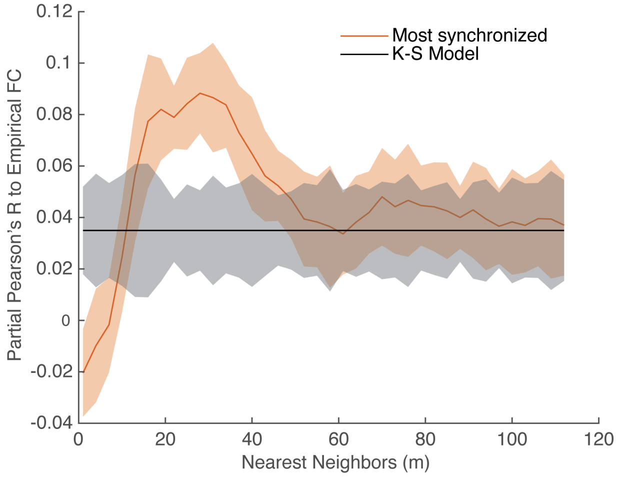


FIG. 1. **Partial correlation to empirical structural connectivity does not explain improved model performance.** The correlation to empirical functional connectivity (FC), controlling for the correlation to empirical structural connectivity is shown for runs of the model across all values of $m$ for $k = 280$. The partial correlation to empirical FC of the K-S model is also shown in black. Clouds indicate a 95% confidence interval. The improved empirical fit seen at and around $m = 19$ is still present (two independent sample t-test, 398 degrees of freedom: $t = 44.72$, $p = 2.58 \times 10^{-157}$), indicating that it cannot be explained by greater similarity to the underlying structural connectivity.

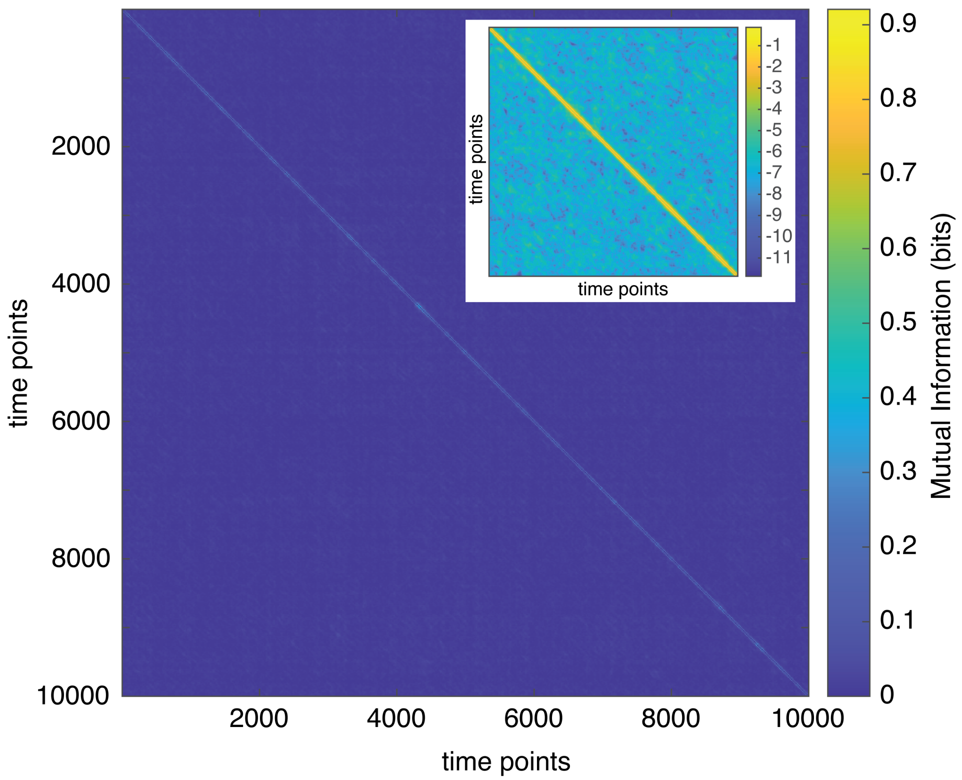


FIG. 2. **Recurrence analysis reveals a large space of selected structural subgraphs.** The mutual information between selected subgraphs was computed for every pair of time points in one run of the model choosing most synchronized neighbors with $m = 19$. The plot shows a sample of the recurrence of 10,000 time points. The matrix is very flat, indicating that selected subgraphs rarely recur. The bright diagonal indicates a slow decay of the current subgraph. To visualize in more detail, the inset shows the mutual information between every pair of the first 2000 time points with a $\log_2$ passed over it to highlight the existing structure.
